## Supplementary Material Jeong, Gonzalez-Fernandez et al for "Micro-Scale Control of Oligodendrocyte Morphology and Myelination by the Intellectual Disability-Linked Protein Acyltransferase ZDHHC9"

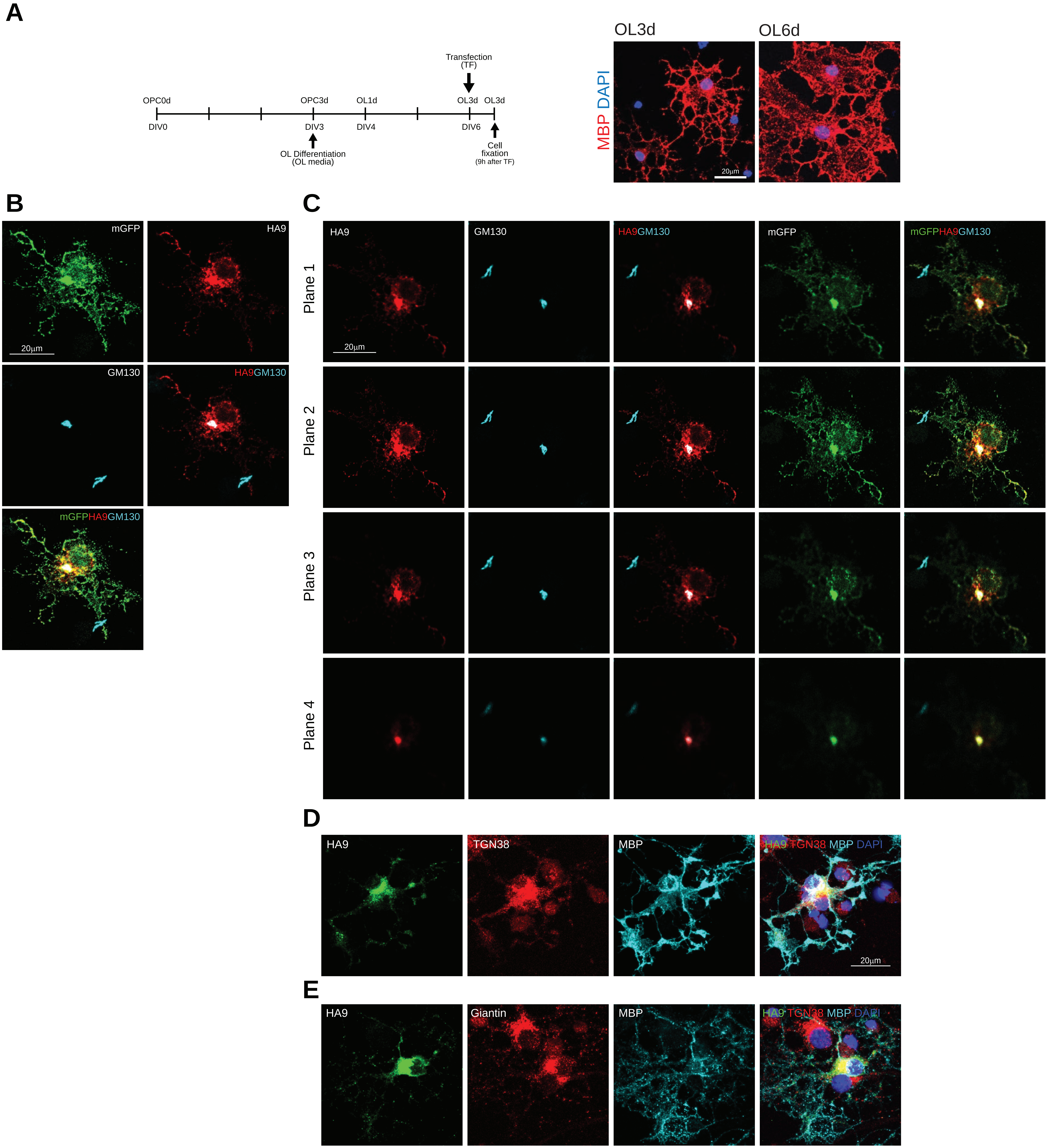


Jeong, Gonzalez-Fernandez et al., Figure S1

**Figure S1: In OL cell bodies, ZDHHC9 localizes to somatic Golgi.** ***A:*** *Left:* Experimental timeline for OL transfection experiments (duplicated from Fig 2A). *Center, right*: Images (maximum intensity projection of confocal z-stacks of cultured OLs after 3 days' incubation in differentiation medium ('OL3D'). Note extensively ramified morphology and MBP positivity, both of which are even more apparent after 6 days' differentiation ('OL6D'). ***B:*** Images of cultured OL (maximum intensity projection of confocal z-slices) transfected to express mGFP and HA-ZDHHC9 (timeline as in Fig 2A) and immunostained as indicated. ***C:*** Images of individual z-slices (planes 1-4) of the cell from *A* confirms colocalization of HA-ZDHHC9WT with Golgi marker GM130. ***D:*** Representative image of OL transfected to express HA-ZDHHC9WT (HA-9) and immunostained to detect HA, MBP and Golgi marker TGN38. ***E:*** As *D*, but for OL immunostained to detect HA, MBP and Golgi marker Giantin.

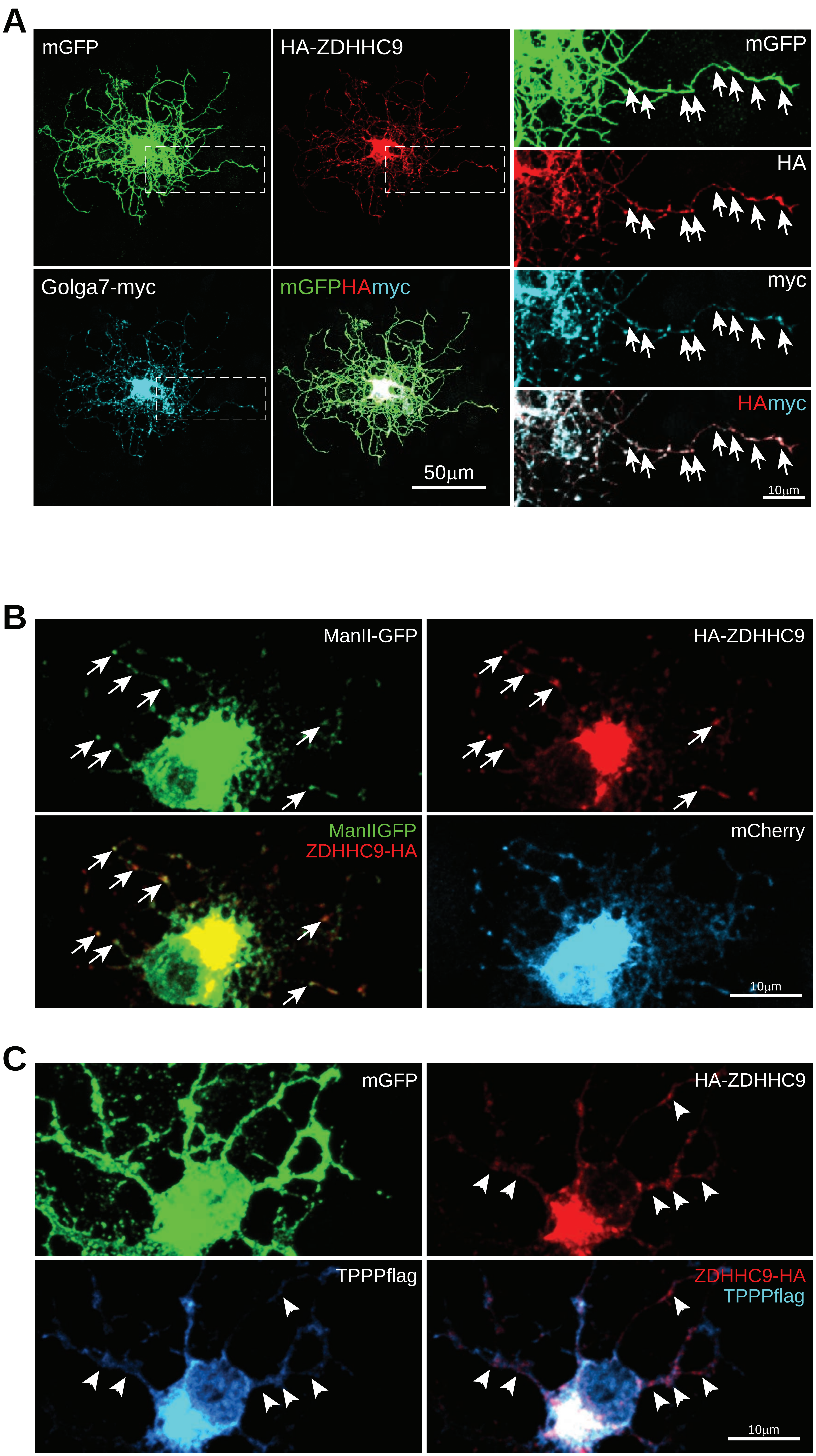


Jeong, Gonzalez-Fernandez et al., Figure S2

**Figure S2: ZDHHC9-positive puncta in OL processes are Golgi outposts/satellites** *A:* Representative image of mature OL, transfected with the indicated cDNAs (timeline as in Fig.7A) and immunostained with the indicated antibodies. HA-ZDHHC9 WT colocalizes extensively with Golga7-myc in puncta in OL processes (arrows in zoomed images). ***B:*** As *A***,** but for OLs transfected to express HA-ZDHHC9 WT, Golgi outpost marker ManII-GFP and mCherry. HA-ZDHHC9 colocalizes extensively with ManII-GFP in OL processes (arrows). *C:* As ***A***, but for OLs transfected to express HA-ZDHHC9 WT, Golgi outpost marker TPPP-Flag and mGFP. HA-ZDHHC9 WT also colocalizes with TPPP-Flag in OL processes (arrows).

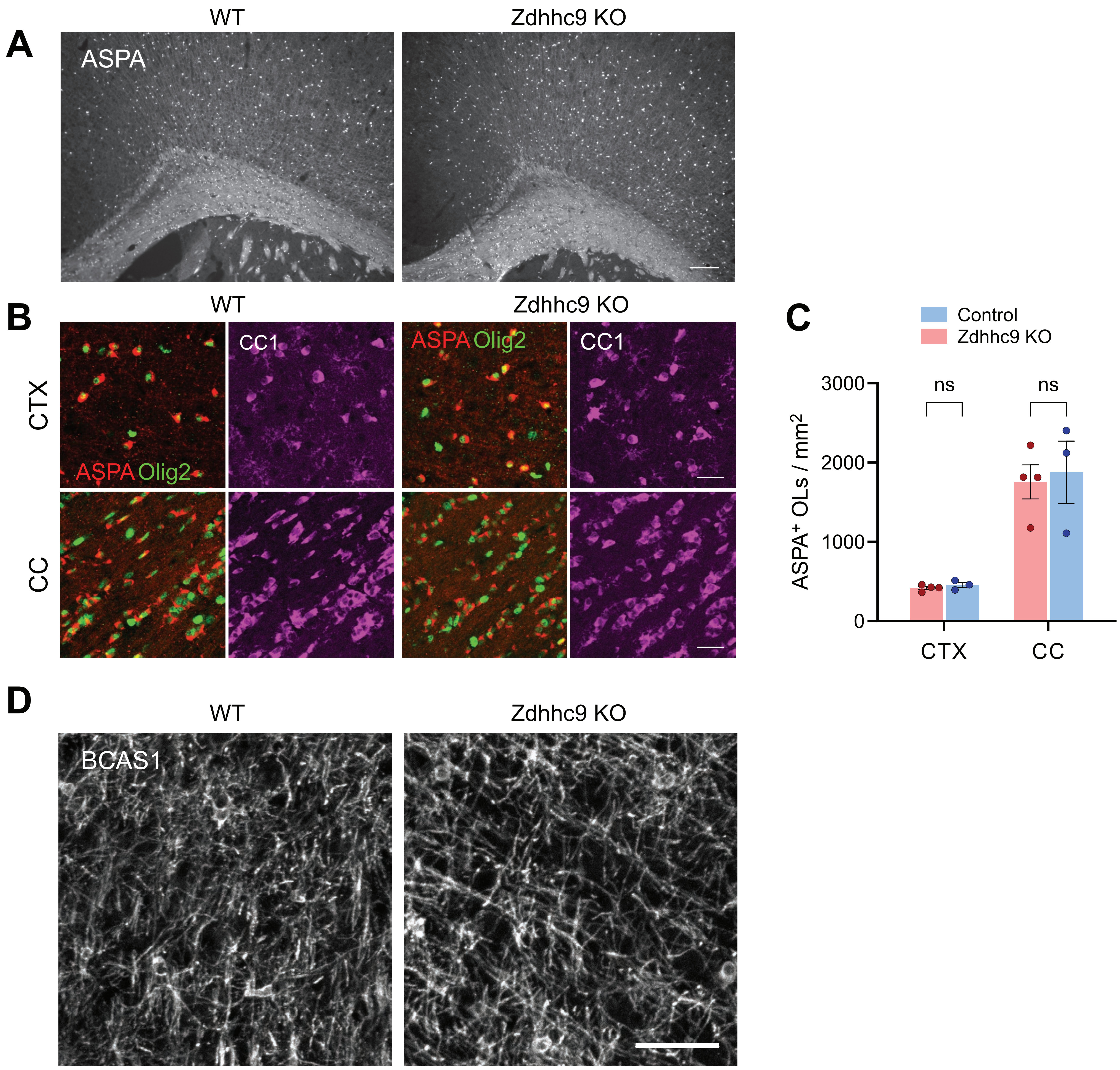


**Figure S3: No alteration of oligodendrocyte density in *Zdhhc9* KO mice. *A:*** Fluorescent images of ASPA$^{+}$ OLs in WT and *Zdhhc9* KO mice. Scale bar: 500 µm. ***B:*** Confocal images of ASPA, CC1 and Olig2 in the CTX and CC. Scale bar 25 µm. ***C:*** Quantified ASPA$^{+}$ OL denstities. Data are mean ± SEM. Students t-test. ns: non-significant. P56 WT (n=4) and *Zdhhc9* KO (n=3) male mice.***D:*** Representative confocal images of BCAS1 in the CTX of 3-week-old WT and *Zdhhc9* KO mice. Scale bar: 50 µm.

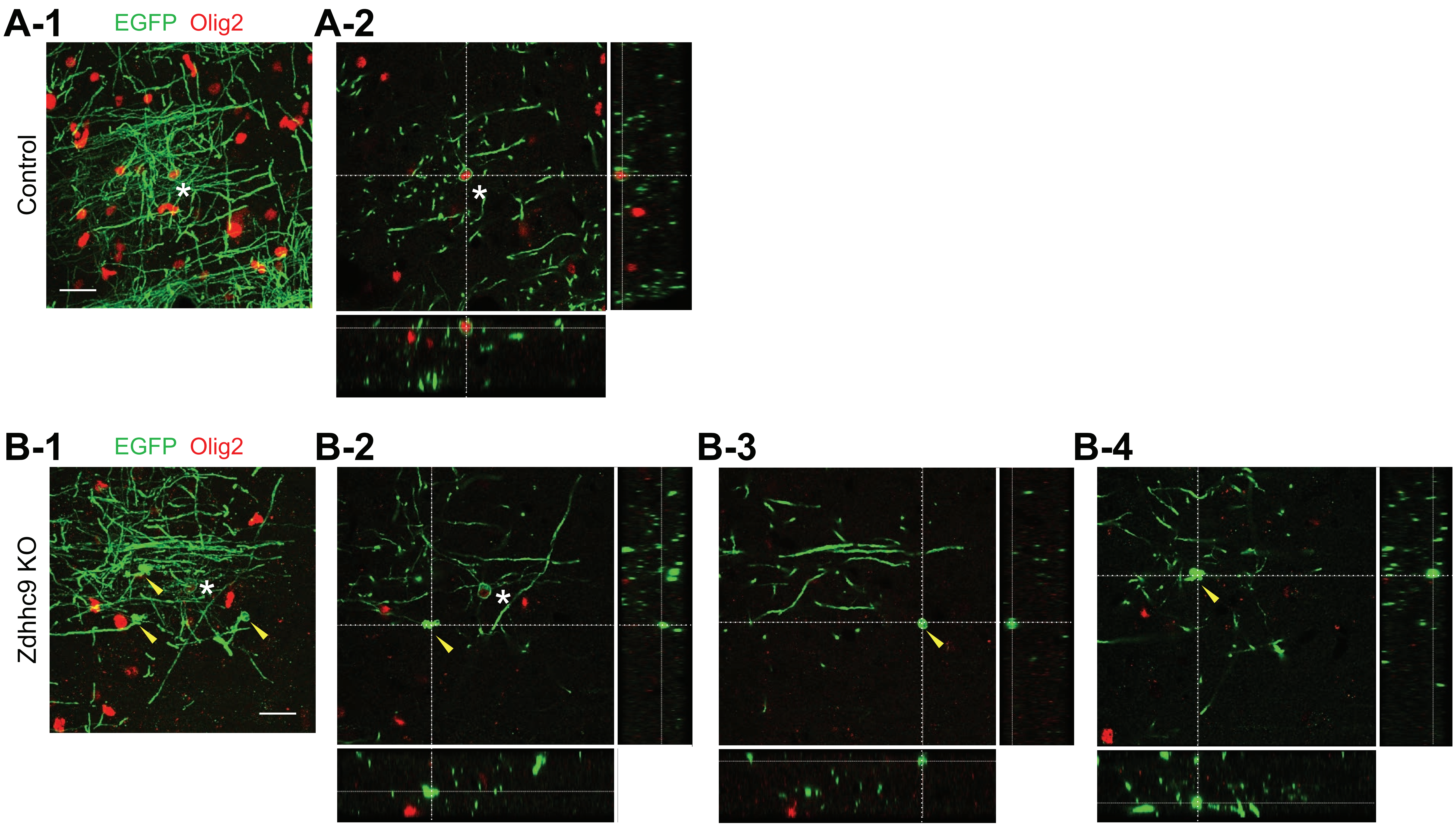


**Figure S4: Spheroid-like abnormal structures in *Zdhhc9* KO mice are distinct from Olig2+ oligodendrocyte cell bodies.** Representative stacked confocal images of EGFP$^{+}$ OLs and Olig2 in control (A-1) and *Zdhhc9* KO (B-1) mice. Orthogonal views of the confocal images shown in A-1 (A-2) or in B-1 (B-2, B-3 and B-4). Scale bars: 20 $\mu$m. Asterisks denote Olig2$^{+}$ cell bodies. Yellow arrowheads indicate abnormal swellings.

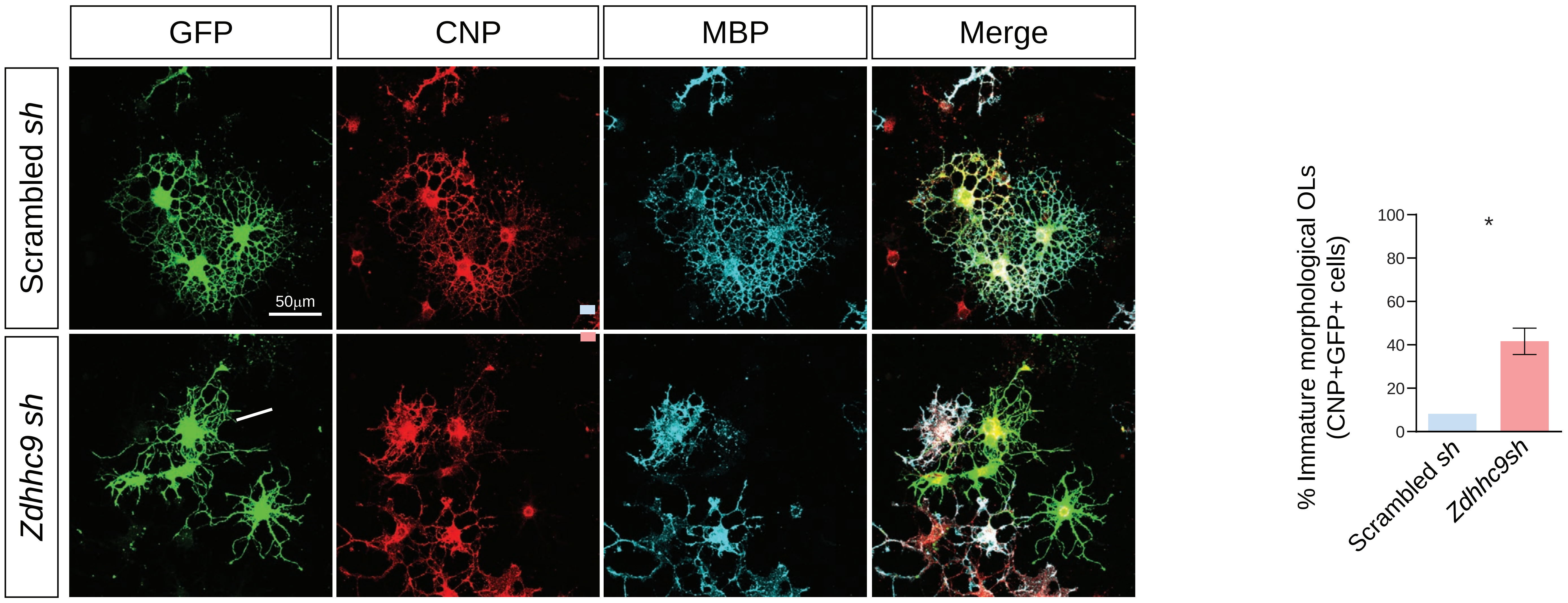


**Figure S5: Zdhhc9 knockdown causes morphological immaturity of committed OLs.** ***A:*** Images of cultured OLs immunostained with the indicated antibodies. Quantified data from *A* confirm that *Zdhhc9* knockdown increases the percentage of immature (CNP$^+$, MBP$^-$) virally infected OLs, n=15 fields of view per condition from 2 coverslips per condition, from a single culture.

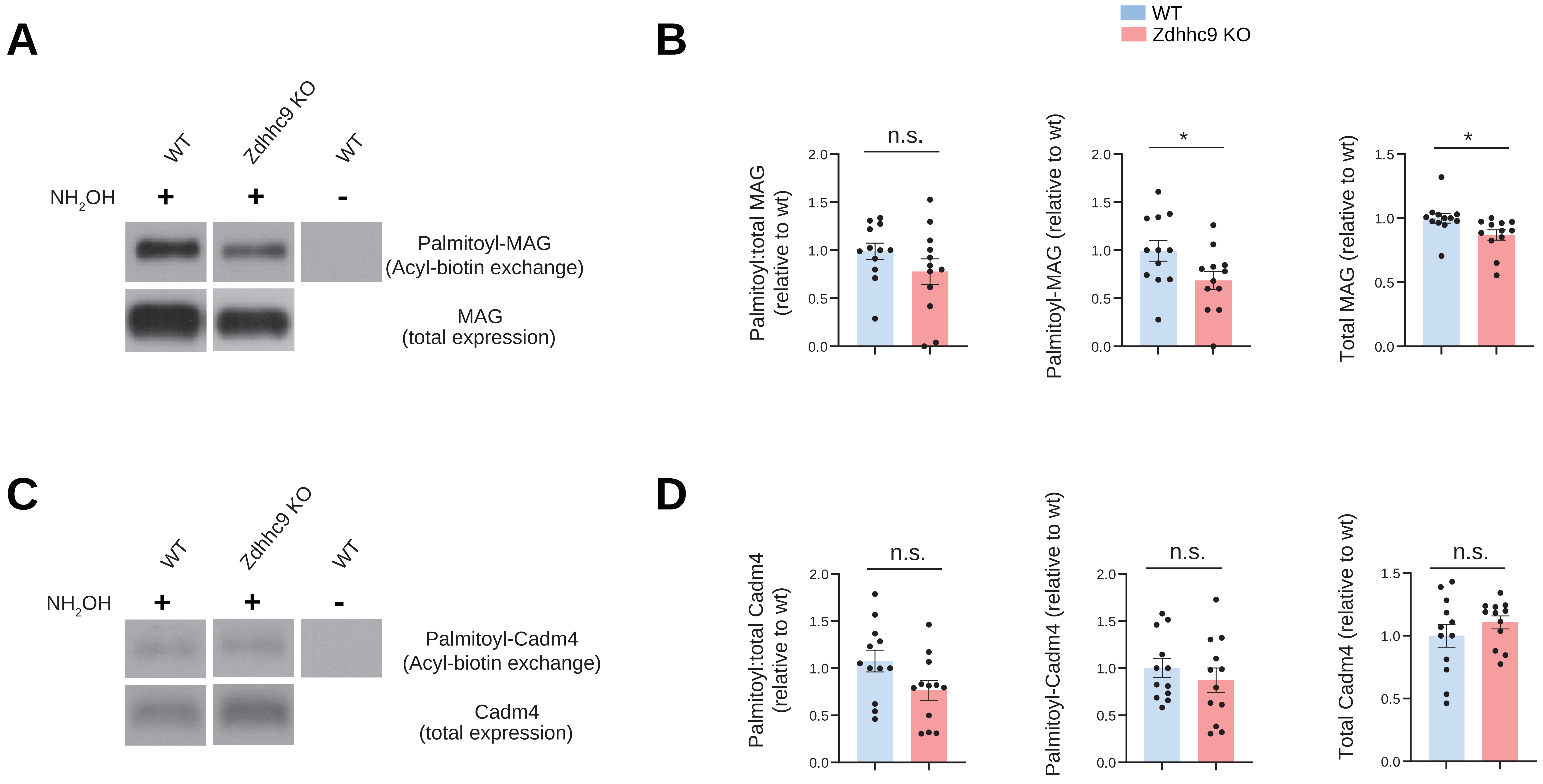


**Figure S6: Zdhhc9 loss impacts the palmitoyl-protein MAG but not Cadm4. *A:*** Western blots to detect MAG in total lysates and ABE fractions from forebrain WM (CC and striatum) from mice of the indicated genotype. ***B:*** Quantified data from *A* confirm that Zdhhc9 loss significantly reduces palmitoyl and total levels of MAG, but palmitoyl:total level is not significantly affected. ***C:*** As *A*, but blotted to detect Cadm4. D: Quantified data from *C* confirm that Zdhhc9 loss does not significantly affect palmitoyl-, total or palmitoyl:total levels of Cadm4. *:$p<0.05$, n.s.; not significant, unpaired t-test, N=12 per genotype.
